# Select DYRK1A Inhibitors Enhance Both Proliferation and Differentiation in Human Pancreatic Beta Cells

**DOI:** 10.1101/2024.05.17.594179

**Authors:** Peng Wang, Olivia Wood, Lauryn Choleva, Hongtao Liu, Esra Karakose, Luca Lambertini, Aidan Pillard, Vickie Wu, Adolfo Garcia-Ocana, Donald K. Scott, Kunal Kumar, Robert J. DeVita, Andrew F. Stewart

## Abstract

The small molecule DYRK1A inhibitor, harmine, induces human beta cell proliferation, expands beta cell mass, enhances expression of beta cell phenotypic genes, and improves human beta cell function i*n vitro* and *in vivo*. It is unknown whether the “pro-differentiation effect” is a DYRK1A inhibitor class-wide effect. Here we compare multiple commonly studied DYRK1A inhibitors. Harmine, 2-2c and 5-IT increase expression of PDX1, MAFA, NKX6.1, SLC2A2, PCSK1, MAFB, SIX2, SLC2A2, SLC30A8, ENTPD3 in normal and T2D human islets. Unexpectedly, GNF4877, CC-401, INDY, CC-401 and Leucettine fail to induce expression of these essential beta cell molecules. Remarkably, the pro-differentiation effect is independent of DYRK1A inhibition: although silencing DYRK1A induces human beta cell proliferation, it has no effect on differentiation; conversely, harmine treatment enhances beta cell differentiation in DYRK1A-silenced islets. A careful screen of multiple DYRK1A inhibitor kinase candidate targets was unable to identify pro-differentiation pathways. Overall, harmine, 2-2c and 5-IT are unique among DYRK1A inhibitors in their ability to enhance both beta cell proliferation and differentiation. While beta cell *proliferation* is mediated by DYRK1A inhibition, *the pro-differentiation* effects of harmine, **2-2c** and 5-IT are distinct, and unexplained in mechanistic terms. These considerations have important implications for DYRK1A inhibitor pharmaceutical development.

## Introduction

Type 1 diabetes (T1D) and type 2 diabetes (T2D) affects 500 million people worldwide, leading to heart disease, stroke, blindness, kidney failure and premature death (1). Both T1D and T2D exhibit a marked reduction in the number of insulin-producing pancreatic beta cells (2–5). With this beta cell deficiency in mind, human cadaveric pancreas or islet transplantation has emerged as a promising treatment for patients with T1D (6,7). However, the limited availability of human cadaveric islets restricts large scale use of these approaches. To overcome this limitation, alternative approaches, such as the transplantation of beta cells derived from human embryonic stem cells (hESCs) or human induced pluripotent stem cells (hiPSCs) have been explored (8,9). In parallel, significant advances have been made in the development of drugs that can expand or regenerate the residual functional beta cells in individuals with T1D or T2D, as well as in the expansion of beta cells from cadaveric donors for use in transplantation strategies (10–26). Despite these advances, there remains a large unmet need for scalable and affordable pharmacologic approaches to induce regeneration and expansion of the remaining beta cells in people with diabetes.

With this background, several groups have demonstrated that small molecule inhibitors of the enzyme dual tyrosine-regulated kinase 1A (DYRK1A) are able to induce beta cells in adult human cadaveric islets to proliferate *in vitro* (10–27) (**Fig. 1**). Even higher rates of proliferation can be achieved by combining DYRK1A inhibitors with Transforming Growth Factor-β inhibitors (TGF-βI’s) or Glucagon-Like Peptide-1 Receptor agonists (GLP1RAs) *in vitro* (11,12). These events also translate to the *in vivo* setting: for example, harmine, alone or in combination with exendin-4 markedly (300% and 700%, respectively) enhances beta cell mass in human islets transplanted into euglycemic and streptozotocin (STZ)-diabetic immunodeficient mice (16,17).

**Figure 1.**
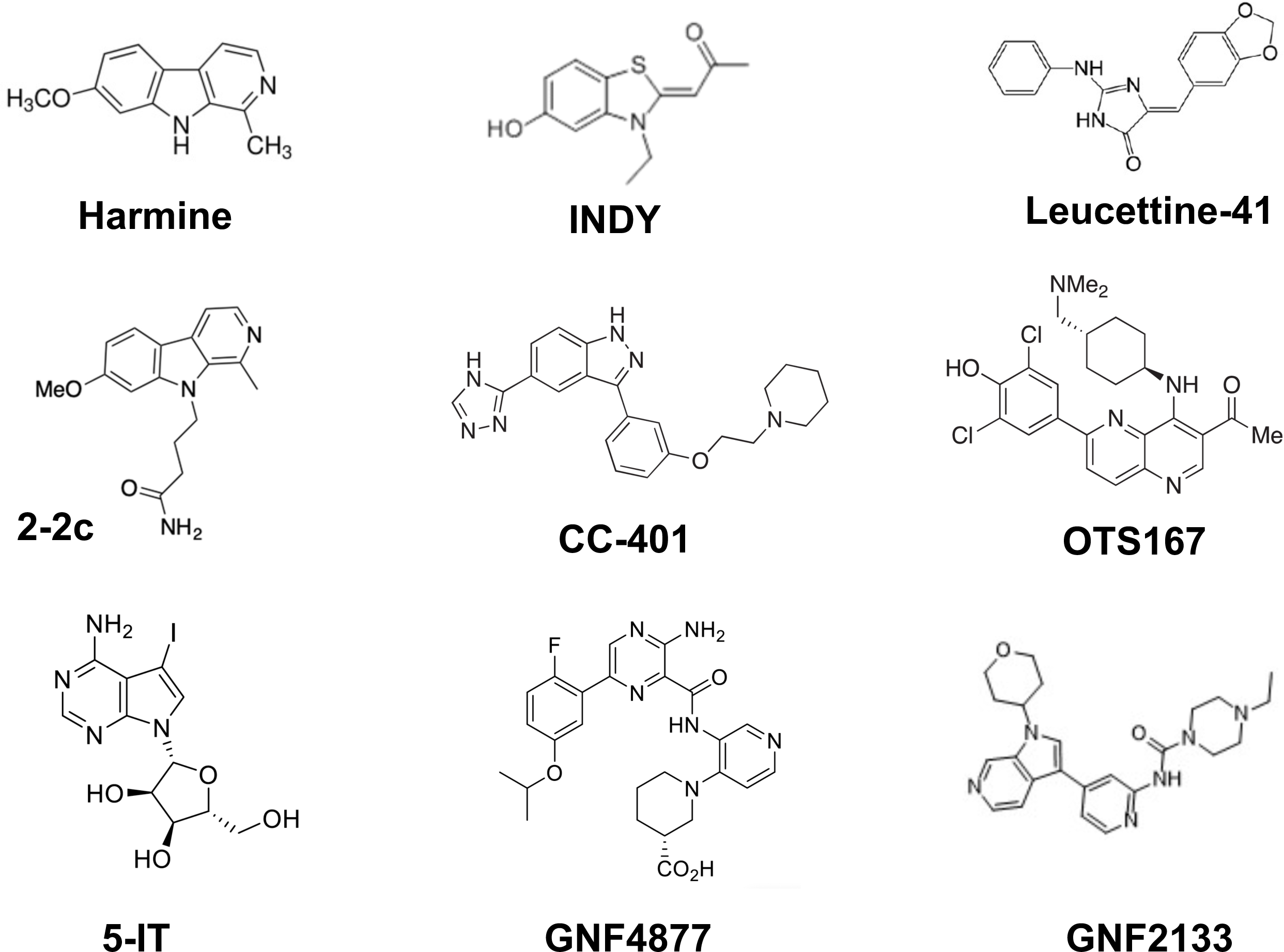
Examples Of Commonly Studied Small Molecule Human Beta Cell Mitogenic DYRK1A Inhibitors. Harmine, INDY, Leucettine-41, 2-2c, CC-401, 5-IT, and GNF4877 used in this study.

It is clear that DYRK1A inhibition is essential for the proliferative effects of harmine and other DYRK1A inhibitors, since silencing DYRK1A in human beta cells fully reproduces the harmine-induced proliferation (10–15,21). Several mechanisms have been proposed to explain how DYRK1A inhibition leads to human beta cell proliferation including inhibition of phosphorylation of: 1) LIN52, an essential component of the cell cycle repressive DREAM complex (14); 2) NFAT transcription factors that transcriptionally regulate cell cycle targets (10,14,28,29); 3) the cell cycle inhibitor, p27^KIP2^, accelerating its degradation (22,23); and, 4) the D-cyclin family, leading to their stabilization and resultant cell cycle activation (22,23). Importantly, all of the molecules in **Fig. 1** are believed to drive proliferation in this manner.

Counterintuitively, perhaps, use of the DYRK1A inhibitor, harmine, alone or together with GLP1 also enhances the differentiation and function of human beta cells. This is evidenced by augmented expression of key beta cell lineage-defining transcription factors, such as PDX1, MAFA, MAFB, NKX6.1, as well as mRNAs and proteins essential for mature beta cell function, exemplified by the beta cell glucose transporter GLUT2, encoded by *SLC2A2,* and the insulin prohormone convertase-1, encoded by *PCSK1* (12). These molecular events lead to enhanced glucose-stimulated insulin secretion (GSIS) from human islets isolated from normal pancreas organ donors, and also from donors with T2D (12). These pro-differentiation effects events also translate to the *in vivo* setting: for example, treatment with harmine, alone or in combination with exendin-4, in (STZ)-diabetic immunodeficient mice transplanted with a minimal mass of human islets, rapidly (days) normalizes blood glucose, and leads to a four-fold increase circulating increases human insulin concentrations (16,17). Thus, harmine is able to simultaneously drive two mechanistically distinct but therapeutically important cellular programs, beta cell proliferation and differentiation, which together make harmine a potentially ideal potential candidate for diabetes treatment.

We assumed that the pro-differentiation effects of harmine also occur in response to *all* small molecule DYRK1A inhibitors. Surprisingly, in the course of the studies described below, we observed that not all DYRK1A inhibitors display this pro-differentiation, function-enhancing effect of harmine: in fact, most do not. We were also surprised to observe that although silencing DYRK1A in human beta cells drives beta cell proliferation, it has no effect on beta cell differentiation. In this report, we explore the mechanisms responsible for the pro-differentiation effect of harmine.

An additional concept is important at the outset. It is well established that no small molecule DYRK1A inhibitor in **Fig. 1** is completely selective for DYRK1A: all DYRK1A inhibitors inhibit other kinases. For example, in conventional kinome-wide screens, GNF4877 interacts with 254 of 468 human kinase (15,24). 5-Iodo-tubericidin (5-IT) is also promiscuous and non-selective, inhibiting 102 of 468 human kinases (15,21). In contrast, harmine and compound **2-2c** are remarkably “clean” or selective, inhibiting only 9 and 8, respectively, of 468 human kinases (15). These include DYRK1A, DYRK1B, DYRK2, and the CDC-like kinases, CLK1, CLK2. In the studies reported below, we pursue a candidate approach to explore these kinases and other possible targets as mediators of the beneficial pro-differentiation effects of harmine.

Finally, a comment on semantics. We have grouped the molecules we chose to study into “beta cell transcription factors” and beta cell “differentiation markers”. We use the term “transcription factor” in its usual sense, referring to PDX1, MAFA, MAFB, NKX6.1, NeuroD1, NKX2.2, PAX4, SIX2, SIX3, NGN3, all well characterized transcription factors essential for early and/or late phases of human beta cell development. In contrast, we use “differentiation factors” advisedly. Some members of this group are critical to human beta cell function, exemplified by SLC2A2, encoding GLUT2, PCSK1 encoding prohormone convertase-1, SLC30A8 encoding the ZnT8 zinc transporter, others have less well defined functions yet are associated with mature beta cell function, exemplified by ENTPD3, which encodes NTPDase-3 or Ectonucleoside Triphosphate Dihydrolase-3, a highly specific surface molecule present on human beta cells and a smaller number of somatostatin-expressing delta cells. We consider these molecules as broad descriptors of the human beta cell phenotype for the purposes of the study.

## Results

### Beta Cell Transcription Factors and Differentiation Markers Are Selectively Induced By Harmine, 5-IT, and 2-2c, But Not Other DYRK1A Inhibitors

To understand the mechanisms underlying the beneficial, pro-differentiation effects of harmine and other DYRK1A inhibitors, we explored the ability of harmine, 5-IT, GNF4877, INDY, Leucettine-41, and CC401 to induce expression, assessed by qPCR in whole human islets, of insulin (INS), beta cell-requisite transcription factors, including PDX1, NKX6.1, MAFA, MAFB, NEUROD and others, as well as other markers of beta cell differentiation, including SLC2A2, GLP1R, PCSK1, SLC30A8, and ENTPD3 (**Figs. 2,3**). We selected doses of the DYRK1A inhibitors previously reported by ourselves and others to yield maximal proliferation rates (10–23). 0.1% DMSO, the diluent for all compounds, was used as a control. Dispersed human islets were treated for 96 hours, and gene expression was analyzed by qPCR. We anticipated that all DYRK1A inhibitors would have similar pro-differentiation effects on human islets.

**Figure 2.**
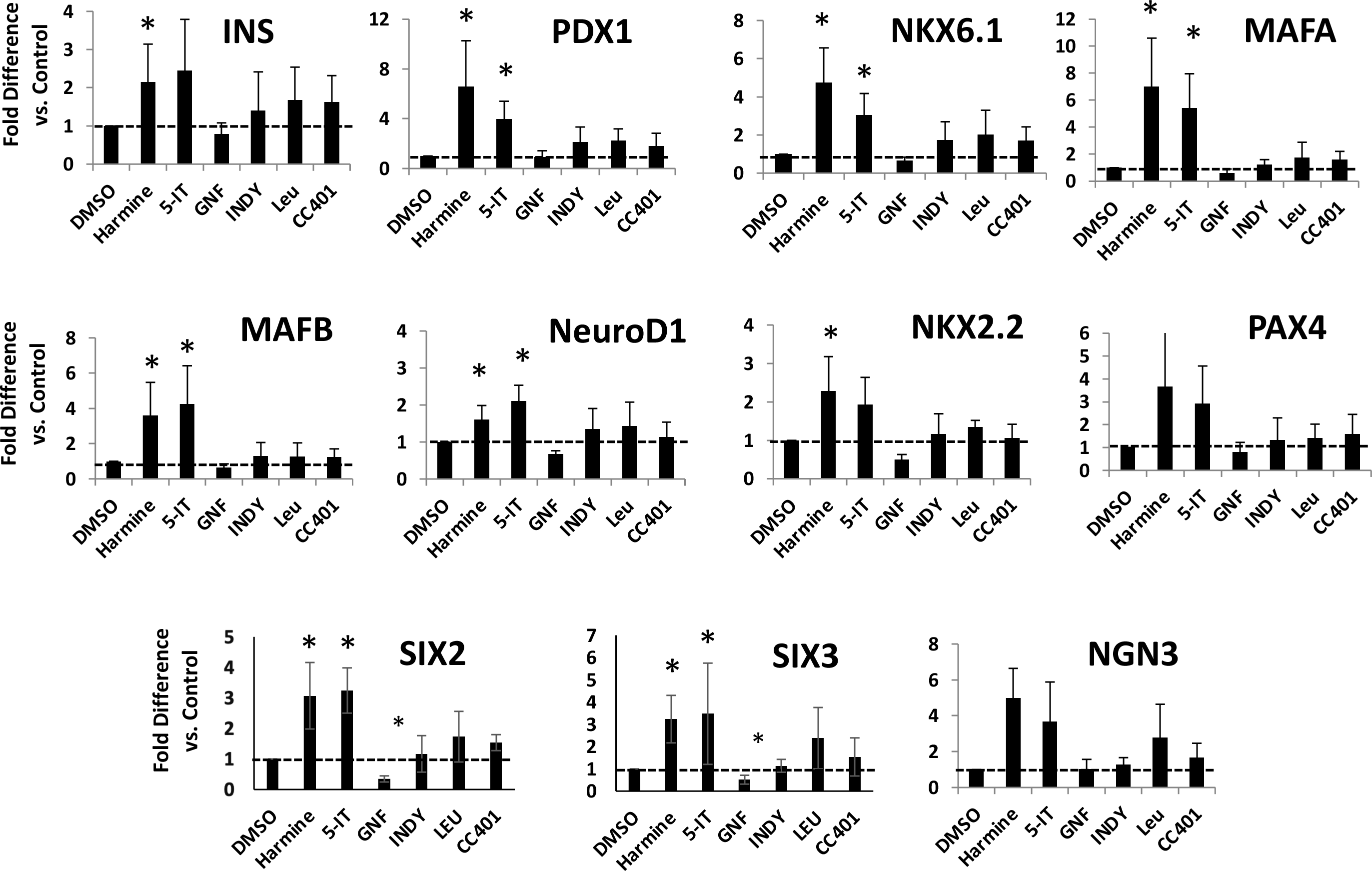
Human Beta Cell Transcription Factors Are Induced By Harmine and 5-IT. Dispersed human islets were treated with DMSO as a control, as well as Harmine, 5-IT, GNF4877, INDY, Leucettine-41, and CC401 at specific concentrations for 4 days, dose chosen based on previous publications demonstrating their highest proliferation rates. The expression of insulin and other major beta cell transcriptional factors was assessed using qPCR. The dotted line highlights the value of the DMSO control. *P <0.05 and **P <0.01 versus control by paired two-tailed t-test. Representative of experiments in five human islet preparations

**Figure 3.**
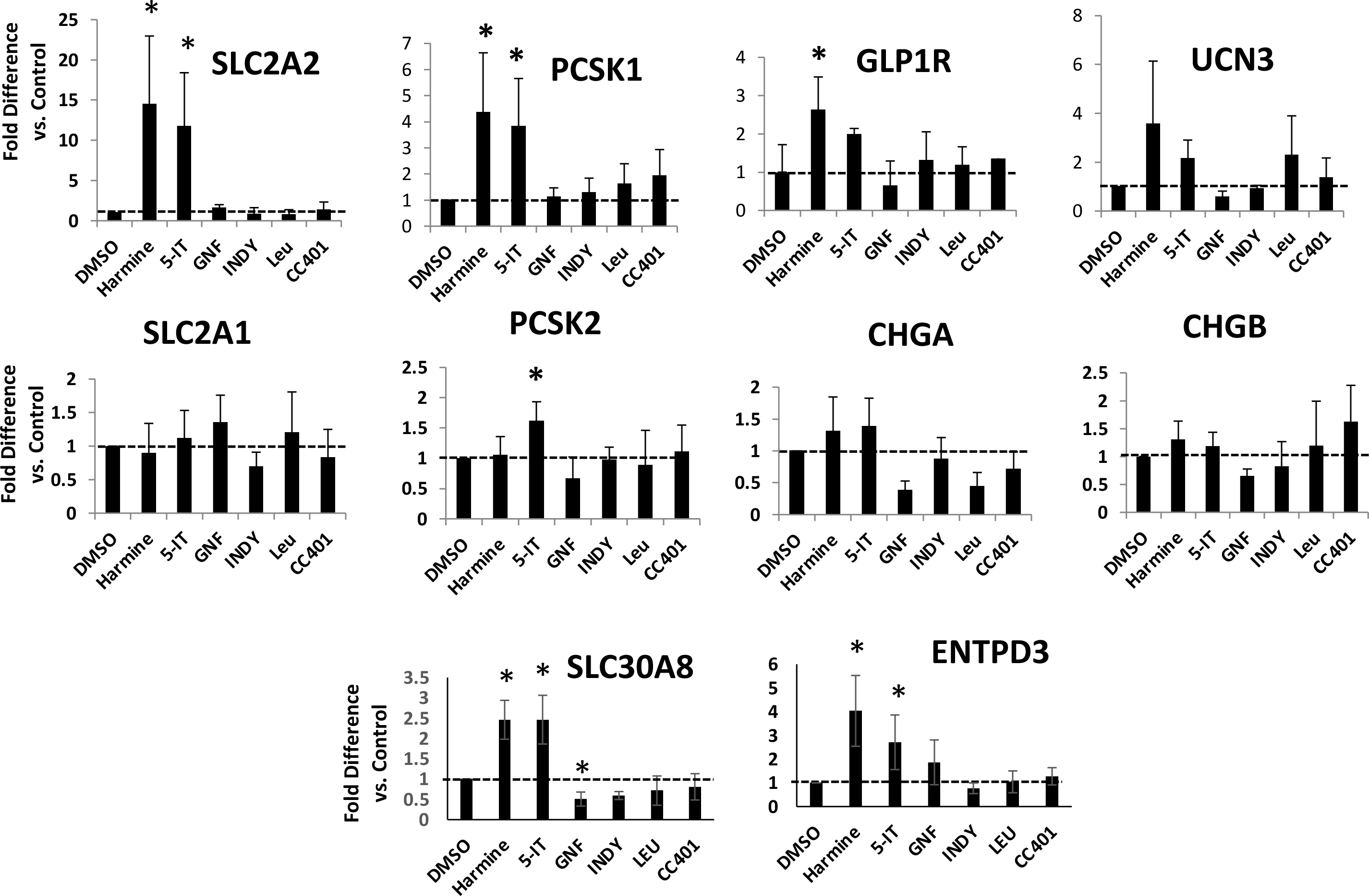
Human Beta Cell Differentiation Markers Are Induced By Harmine and 5-IT. Dispersed human islets were treated with DMSO as a control, as well as Harmine, 5-IT, GNF4877, INDY, Leucettine-41 and CC401for 4 days as in Figure 2. The expression of beta cell differentiation markers was assessed using qPCR. The dotted line highlights the value of the DMSO control. *P <0.05 and **P <0.01 versus control by paired two-tailed t-test. Representative of experiments in five human islet preparations

In contrast to our expectations, we observed that among all DYRK1A inhibitors studied, only harmine and 5-IT were able to augment human beta cell differentiation markers and transcription factors. We also studied a novel more potent harmine-based DYRK1A inhibitor called compound 2-2c **(Suppl. Fig 1)** (15). Since 2-2c is 3-fold more potent than harmine at driving human beta cell proliferation (15), we studied it at a 3-fold lower dose than harmine. At a concentration of 3 µM, 2-2c induced the expression of PDX1, NKX6.1, MAFA, and SLC2A2 to a degree similar to 10 µM harmine **(Suppl. Fig 1)**.

Since harmine has previously been reported to increase protein abundance of insulin, GLUT2 and beta cell transcription factors (12) we performed immunocytochemical studies on human islets. **Fig. 4** demonstrates that the increase in mRNA expression induced by harmine and 5-IT was paralleled by an increase protein expression of PDX1, NKX6.1 and ENTPD3. These effects were not observed with the two other DYRK1A inhibitors studied: GNF4877 and INDY.

**Figure 4.**
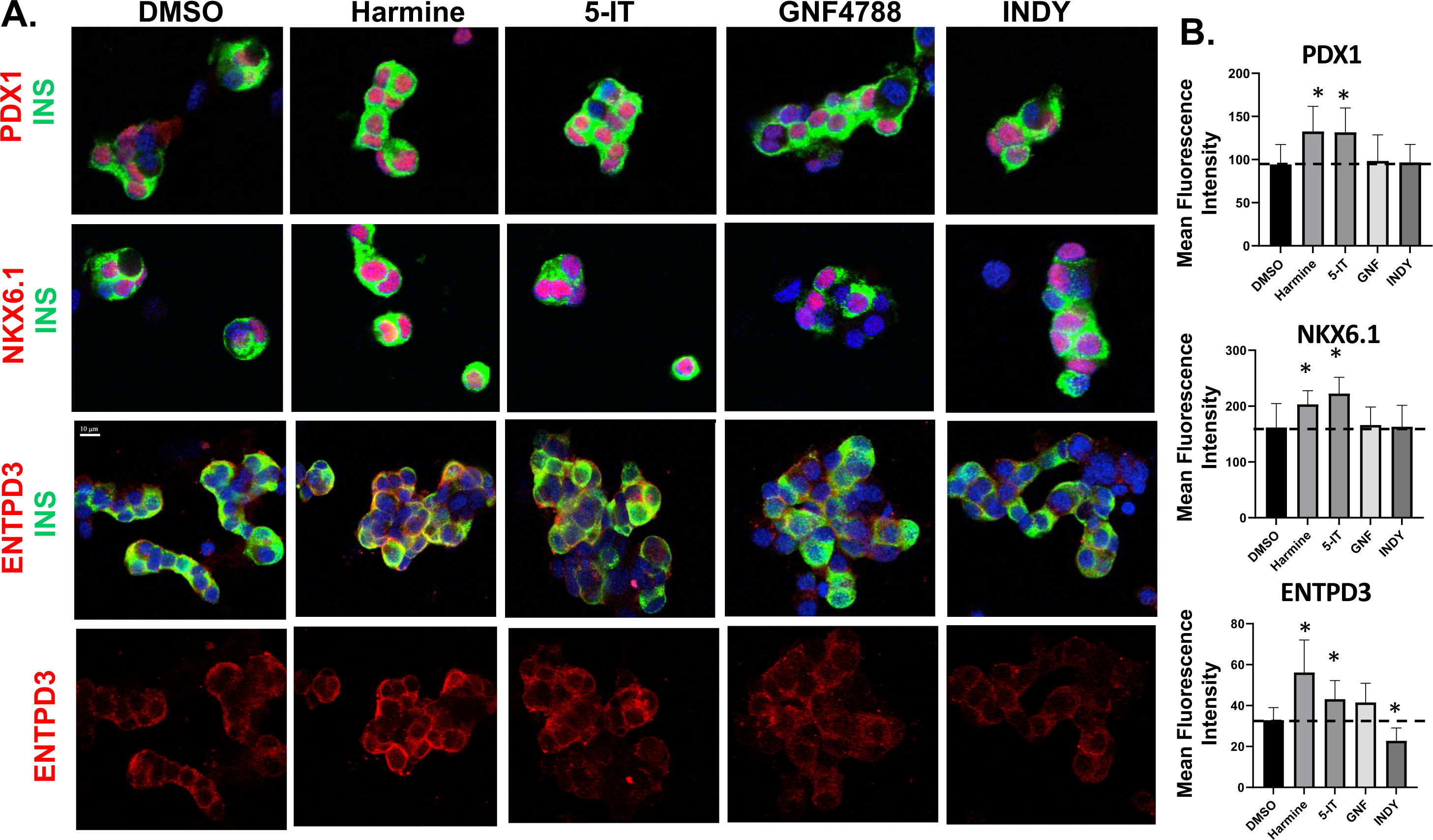
Harmine and 5-IT Induce Greater Increases in PDX1, NKX6.1 and ENTPD3 Protein Abundance Than GNF4877 or INDY. **A**. Examples of confocal imaging of dispersed human islets immunolabelled for INS, PDX1, NKX6.1 and ENTPD3 following treatment with DMSO, Harmine, 5-IT, GNF4877 or INDY for 96 hours. Representative of experiments in five human islet preparations. **B**. The fluorescence intensity of the protein expression quantified by image J. The dotted line highlights the value of the DMSO control. *P <0.05 and **P <0.01 versus control by paired two-tailed t-test. Representative of experiments in five human islet preparations

### GNF4877 Is Non-Selective: It Inhibits Beta Cell Transcription Factors and Interferes with Harmine-Induced Beta Cell Differentiation

Since GNF4877 is a particularly non-selective kinase inhibitor (15,24), we wondered whether it may interfere with cellular processes required for beta cell differentiation. Accordingly, we studied it at multiple concentrations from 0.5µM to 10µM, above and below its optimal beta cell mitogenic concentration, 2 µM (13,15,24). As shown in **Fig. 5**, in contrast to harmine which induced MAFA, NKX6.1, PDX1, GLUT2, PCSK1, and GLP1R expression, GNF4877 at all doses either failed to induce or actually inhibited expression of these key beta cell genes. Remarkably, GNF4877 actually inhibited harmine induced expression of MAFA, NKX6.1 and PDX1. These events are consistent with GNF4877 having non-specific, undesirable, off-target effects on pathways essential for beta cell differentiation. Notably, although 5-IT is also a non-specific DYRK1A inhibitor like GNF4877 (15,21), it actively induces beta cell GLUT2/SLC2A2 expression (**Figs. 2-4**). We suspect that it does not interact adversely with the unknown deleterious targets of GNF4877.

**Figure 5.**
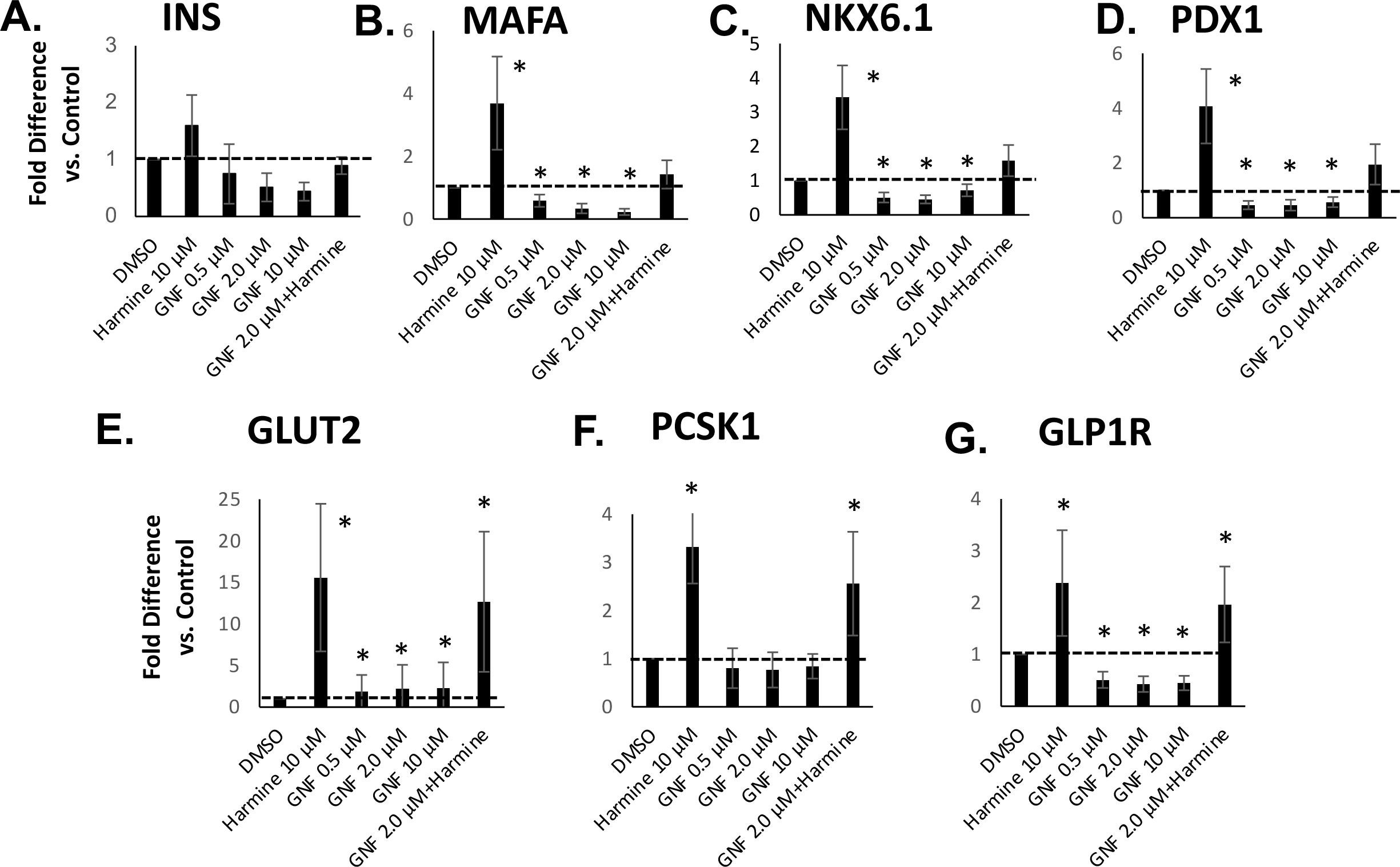
GNF4877 Is Non-Selective: It Inhibits Abundance of Some Transcription Factor and Differentiation Marker mRNAs and Interferes with Harmine Actions. Dispersed human islets were treated with GNF4877 at various concentrations, ranging from 0.5 µM to 10 µM with or without harmine for 96 hours. The expression of insulin and other major beta cell transcriptional factors and differentiation markers was assessed by qPCR. The data demonstrate GNF4877 interfered with the transcription factor expression induced by harmine. The dotted line highlights the value of the DMSO control. *P <0.05 and **P <0.01 versus control by paired two-tailed t-test. Representative of experiments in five human islet preparations.

### DYRK Family Members Do Not Mediate the Expression of Beta Cell Transcription Factors and Differentiation Markers Induced by Harmine, 5-IT, and 2-2c

We next turned to our candidate gene approach. We assumed initially that DYRK1A and DYRK1B might participate in the pro-differentiation effects of harmine, 2-2c and 5-IT, since all share the ability to inhibit DYRK1A and DYRK1B (10–13,15,21,27). Thus, we next queried whether silencing of DYRK1A and DYRK1B in human islets might mimic the pro-differentiation effect of harmine, 2-2c and 5-IT. To assess this possibility, we used a previously described adenovirus (13) encoding shRNAs that effectively silence both DYRK1A and DYRK1B in human islets. **Fig. 6A** demonstrates effective silencing of DYRK1A and DYRK1B in human islets. **Figs. 6C-H** make several important points. First, adenoviral transduction with a scrambled shRNA adenovirus has no effect on differentiation markers. Second, harmine used in the presence of the control shRNA-encoding adenovirus has its usual pro-differentiation effects. Third, transduction of human islets with the shDYRK1A/DYRK1B virus had no effect on human beta cell differentiation markers. And, fourth, simultaneous silencing of DYRK1A and DYRK1B had no deleterious effect on harmine-induced human beta cell differentiation marker gene expression. We conclude that inhibition of DYRK1A and DYRK1B, while essential to human beta cell proliferative effects of DYRK1A inhibitors (12,13), is not involved in the pro-differentiation effects of harmine.

**Figure 6.**
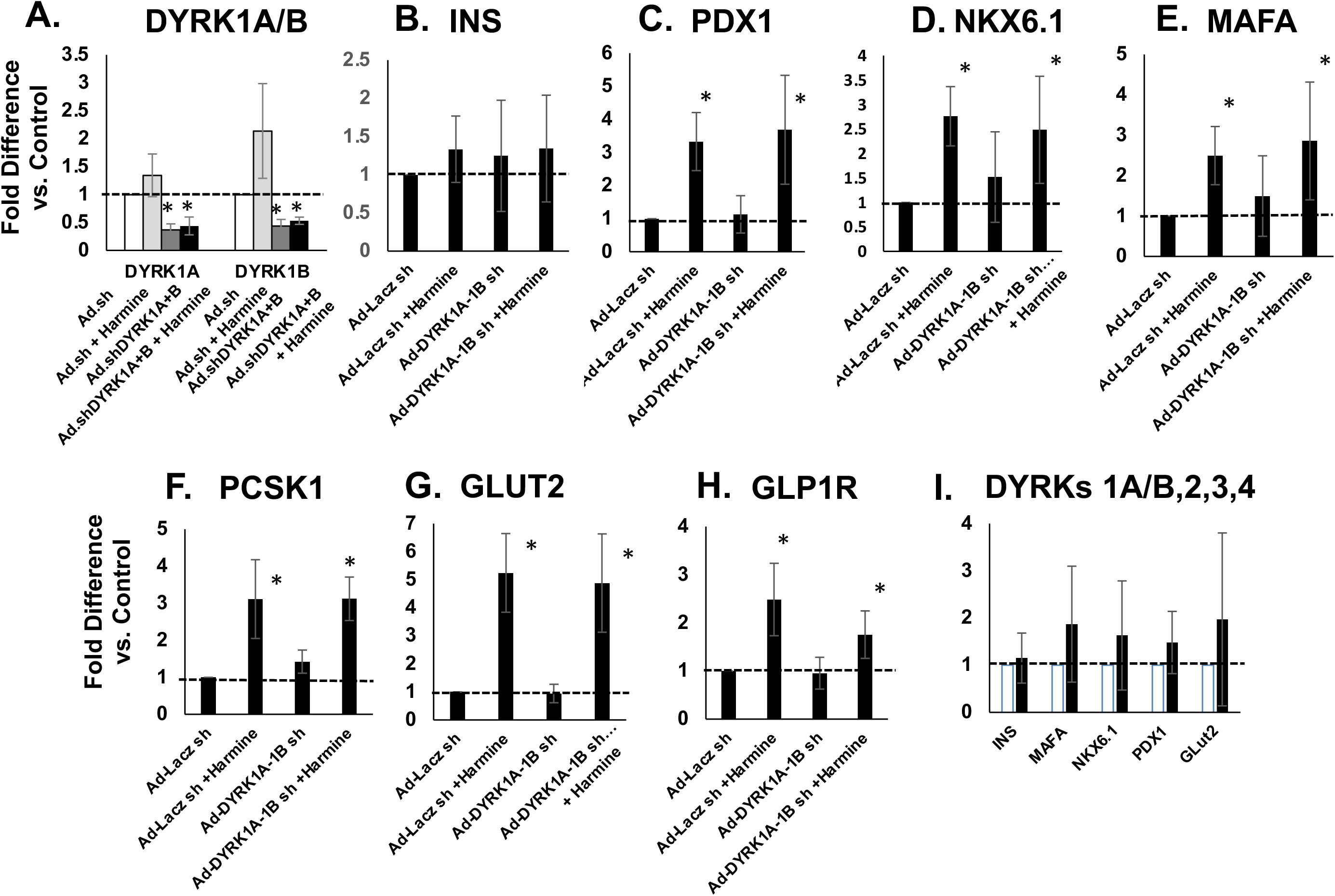
Differentiation Is Not Mediated By The DYRK Family. **A.** qPCR studies in human islets demonstrating that a single combined adenovirus silencing DYRK1A and DYRK1B is effective in reducing DYRK1A and DYRK1B expression (11). **B-H**. For each panel, adenovirus expression of a scrambled shRNA has no effect on target gene abundance (negative control). The same shRNA virus in the presence of harmine increases expression of the target gene (positive control). Silencing DYRK1A/B has no effect on the target gene. Silencing DYRK1A/B in the presence of harmine does not adversely affect harmine-induced differentiation. **I.** Silencing DYRK1A/B,2,3,4 (13) also has no effect on human beta cell differentiation markers. A minimum of five different human islet preparations was used in each panel. The dotted line highlights the control value. *P <0.05 and **P <0.01 versus control by paired two-tailed t-test.

Since most small molecule DYRK1A inhibitors also inhibit activity of the other DYRK family members (13,15,27), we also studied our previously described adenovirus (13) that broadly silences all DYRK family members, including DYRK1A, DYRK1B, DYRK2, DYRK3 and DYRK4. Silencing all DYRK family members did not increase the expression of beta cell transcriptional factor and differentiation markers as showing in **Fig. 6I**. Collectively, these findings indicate that although silencing DYRK1A supports beta cell proliferation, neither DYRK1A nor other DYRK family members appear to participate in human beta cell differentiation.

### The DREAM Complex Does Not Mediate Beta Cell Transcription Factor and Differentiation Marker Expression Induced by Harmine, 5-IT, and 2-2c

As described in the Introduction, the DREAM complex, a key pathway regulating human beta cell proliferation induced by DYRK1A inhibitors, is also a potential candidate mediator of DYRK1A inhibitor-induced differentiation (14). To assess involvement of the DREAM complex in driving human beta cell differentiation, we silenced key DREAM complex target genes encoding the transcription factors, E2F4 and E2F5, in human islets (**Fig. 7A**). While silencing these two genes induces proliferation in human islets (10,13,14), this had no effect on human beta cell differentiation markers.

**Figure 7.**
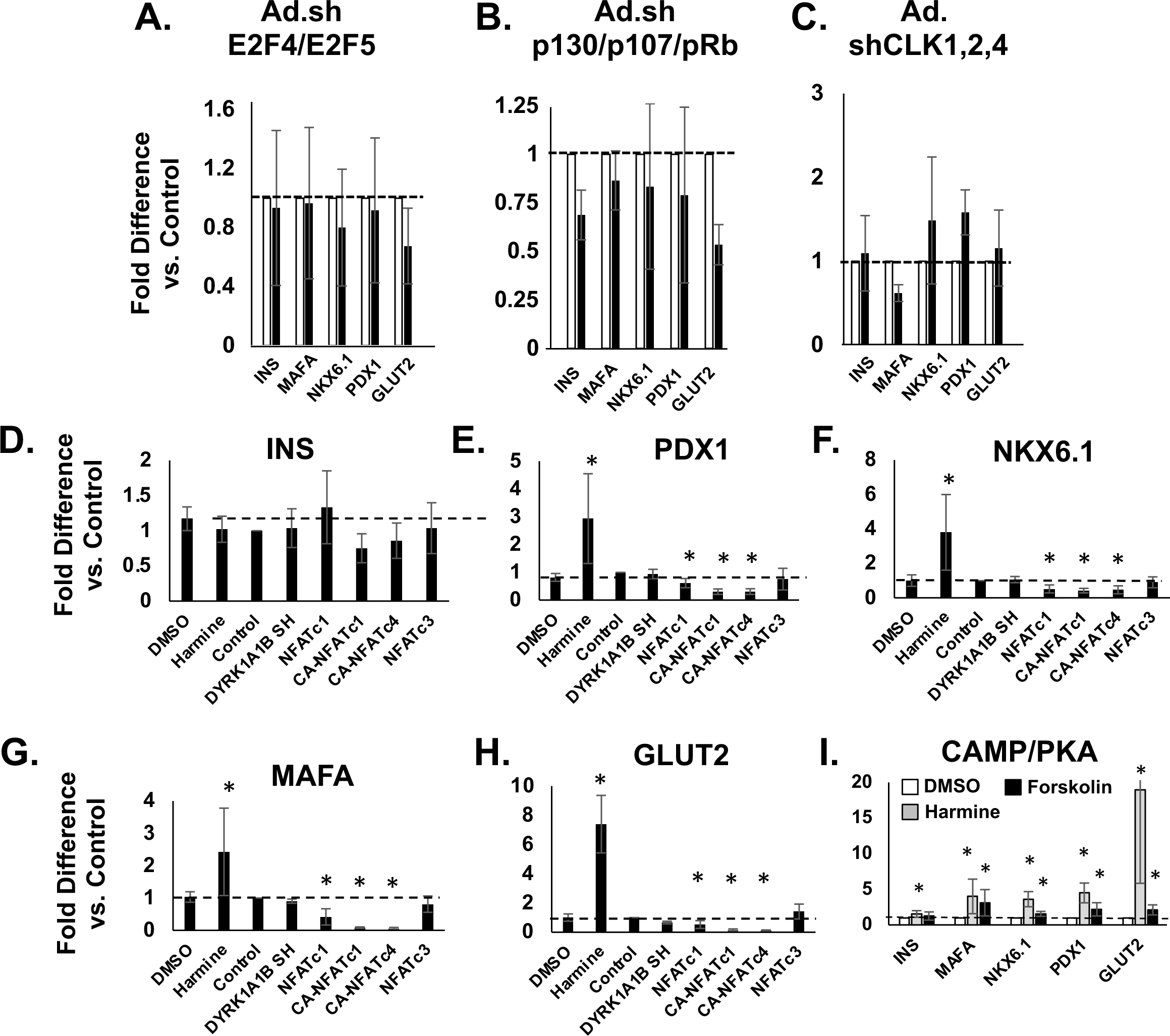
Differentiation Is Not Mediated By The DREAM Complex, CLKs 1,2,4, NFaTs, or cAMP/PKA. **A,B.** qPCR studies of human islets in which key DREAM complex repressive members E2F5 and E2F5 and p107, p107 and pRb were silenced (14) has no effect on human beta cell differentiation markers. **C.** Silencing CLKs 1,2,4 (13) has no effect on expression of target genes. **D-H.** Overexpressing normal NFAT family members, and constitutively active NFAT versions (14) has no effect on human beta cell transcription factor or differentiation marker expression. **I.** Harmine induces differentiation markers, but the activation of cAMP with 10 µM forskolin has modest effects. The dotted line highlights the control value. *P<0.05 and **P<0.01 versus control by paired two-tailed t-test. N=3-5 human islet donors per panel.

In similar studies, we used an adenovirus that silences three other key downstream cell cycle repressive “pocket proteins”, pRb, p107 and p130. While silencing the three pocket proteins with this virus activates human beta cell proliferation (14), this had no effect on human beta cell differentiation (**Fig. 7B**). Collectively, these studies make it unlikely that the DREAM complex is involved in the pro-differentiation effects of harmine, 2-2c and 5-IT.

### The CLK Family Does Not Mediate Beta Cell Transcription Factor and Differentiation Marker Expression Induced by Harmine, 5-IT, and 2-2c

Pursuing the candidate target approach, we next focused on the CLK family using an adenovirus we have employed and validated previously (13) that silences CLKs 1, 2 and 4, the three CDK-like kinases that interact with harmine and 2-2c (**Suppl. Fig. 1**). As be seen in **Fig. 7C**, silencing these three CLKs in human islets had no effect on markers of human beta cell transcription factors or differentiation markers.

### NFAT Family Members and cAMP/PKA Do Not Mediate Beta Cell Transcription Factor and Differentiation Marker Expression Induced by Harmine, 5-IT, and 2-2c

We next focused on NFAT family members as candidate drivers of human beta cell differentiation, since they are well described substrates for DYRK1A phosphorylation, and have been implicated in driving beta cell proliferation in response to harmine and other DYRK1A inhibitors (10,14,28,29). Thus, we overexpressed wild type and constitutively active, nuclear localizing forms of multiple NFAT family members in human islets (14,29) and examined expression of differentiation markers and transcription factors (**Fig. 7D-G**). In contrast to our expectations, while harmine enhanced expression of PDX1, NKX6.1, MAFA and GLUT2 as expected, none of the overexpressed NFAT family members induced expression of these genes, and several actually reduced their expression. Although NFATs are induced to translocate to the nucleus in human beta cells in response to harmine and 2-2c, these observations make it unlikely that they participate in the enhanced differentiation response to harmine and 2-2c.

The cAMP/PKA pathway is required for the synergistic beta cell proliferation induced by the combination of harmine and GLP1 (12), suggesting a possible role for cAMP/PKA signaling as a candidate for the enhanced differentiation induced by harmine alone or in combination with GLP1. Thus, we queried whether the cAMP activator, forskolin, could induce human beta cell differentiation markers (**Fig. 7I**). Here, we found that while harmine had the expected effects, the effects of forskolin were modest and less than harmine treatment. These findings suggest that cAMP/PKA signaling is not a major contributor to the enhanced expression of transcription factor and differentiation markers induced by harmine.

### Harmine, 5-IT, and 2-2c Induce Beta Cell Transcription Factors and Differentiation Markers in Human Islets from People with T2D

Islets from organ donors with T2D express reduced levels of beta cell transcription factors, but these can be increased by exposure to harmine (12). Therapeutically, therefore, it is important to determine whether the beneficial effects of DYRK1A inhibitors translate to islets from people with T2D. **Figures 8A-F** explore this question and demonstrate that harmine and 5-IT have pro-differentiation effects on T2D islets, similar to those observed in normal islets in **Figures 2**, **3**. In contrast, these effects are not observed with GNF4877, INDY, Leucettine or CC401. Since ALDH1A3 and TXNIP are among the putative mediators of beta cell de-differentiation in beta cells in people with T2D (30–33), we also explored the effects on their expression by the DYRK1A inhibitor panel (**Fig. 8 G,H**). Remarkably, harmine 5-IT and the other DYRK1A inhibitors have beneficial effects on ALDH1A3 and TXNIP expression.

**Figure 8.**
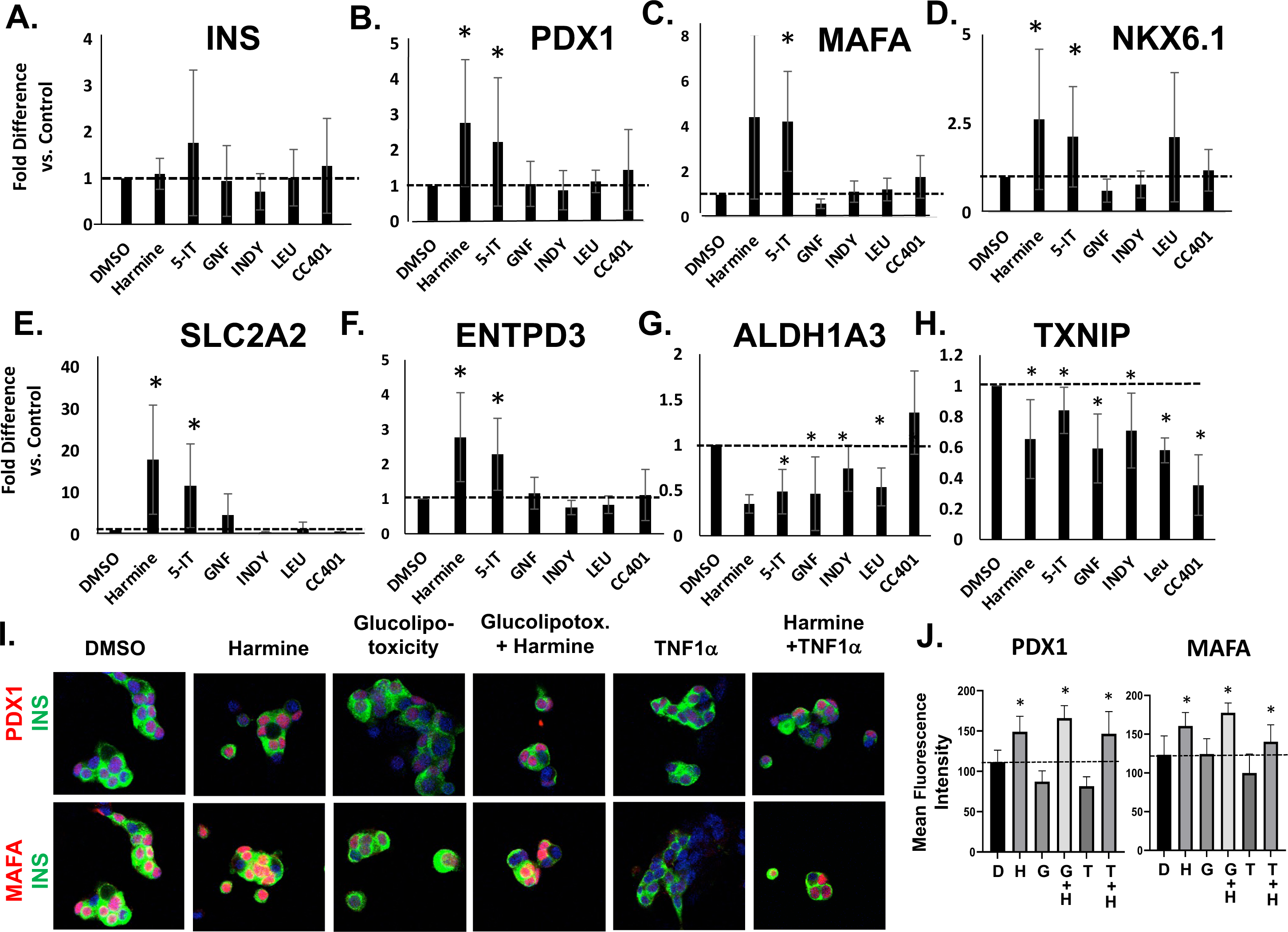
Harmine and 5-IT Induce Beta Cell Differentiation Genes in Islets From T2D Donors. **A-F.** qPCR results in islets from T2D donors in response to the panel of DYRK1A inhibitors. **G-H.** The responses of ALDH1A3 and TXNIP to DYRK1A inhibitors. See text for details. **I.** Immunohistochemical assessment of the effects of DMSO, harmine, glucolipotoxic concentration of glucose (22mM) and palmitate (200μM), and, of TNFα (1000 IU/ml) alone or with harmine (10 μM). **J.** Densitometric analysis of fluorescence intensity of the proteins in panel I. D = DMSO vehicle, H = harmine, G = Glucolipotoxicity, T = TNFα. N = 5 human islet donors per panel. *P<0.05 and **P<0.01 versus control by paired two-tailed t-test.

To determine if the changes observed at the RNA level in T2D islets translate to the protein level, we explored the ability of harmine to protect against glucolipotoxicity-induced (using 20 mM glucose and 250μM palmitate) and cytokine-induced (using TNFα 1000IU) beta cell de- differentiation, as occur in T2D and T1D (**Fig. 8 I,J**). Remarkably, while nuclear expression of PDX1 and MAFA diminished with glucotoxicity and TNFα exposure, harmine appears to have a protective effect in both settings.

## Discussion

Small molecule DYRK1A inhibitors, through their ability to induce human beta cell proliferation *in vitro* and beta cell mass expansion *in vitro* and *in vivo,* have the potential to be transformative for T1D and T2D. They also: 1) enhance human beta cell function, exemplified by glucose-stimulated insulin secretion *in vitro* and *in vivo* (10–27); 2) rapidly reverse diabetes in immunodeficient mice transplanted with a marginal mass of human islets (10,12,16,17); and, 3) augment expression of canonical human beta cell lineage-defining, and function-enabling molecules (12), as illustrated in **Figs 2** and **3**. Mechanistically, it is clear that the proliferation induced by harmine and other DYRK1A inhibitors is mediated by their interference with DYRK1A function, a premise supported by the facts that all DYRK1A inhibitors studied have this effect, and that silencing DYRK1A in human islets is sufficient to activate human beta cell proliferation (10,12,14,21). We, and perhaps others, had assumed that the pro-differentiation properties of harmine also are related to DYRK1A inhibition, and presumably would translate to all small molecule DYRK1A inhibitors. Thus, we were surprised to observe that not all DYRK1A inhibitors are equal, that most members of the class lack this important property. From a therapeutic standpoint, the implications are clear: DYRK1A inhibitors that promote both beta cell proliferation and function, such as harmine, 2 -2c and 5-IT, would be expected to have an advantage for diabetes treatment over other members of the class. From a mechanistic standpoint, however, this phenomenon remains unexplained. One must assume that harmine, 2-2c and 5-IT support differentiation by acting on additional, as yet unidentified, targets unrelated to DYRK1A, as illustrated in **Fig. 9**. Having identified this issue, we next explored potential underlying mechanisms.

**Figure 9.**
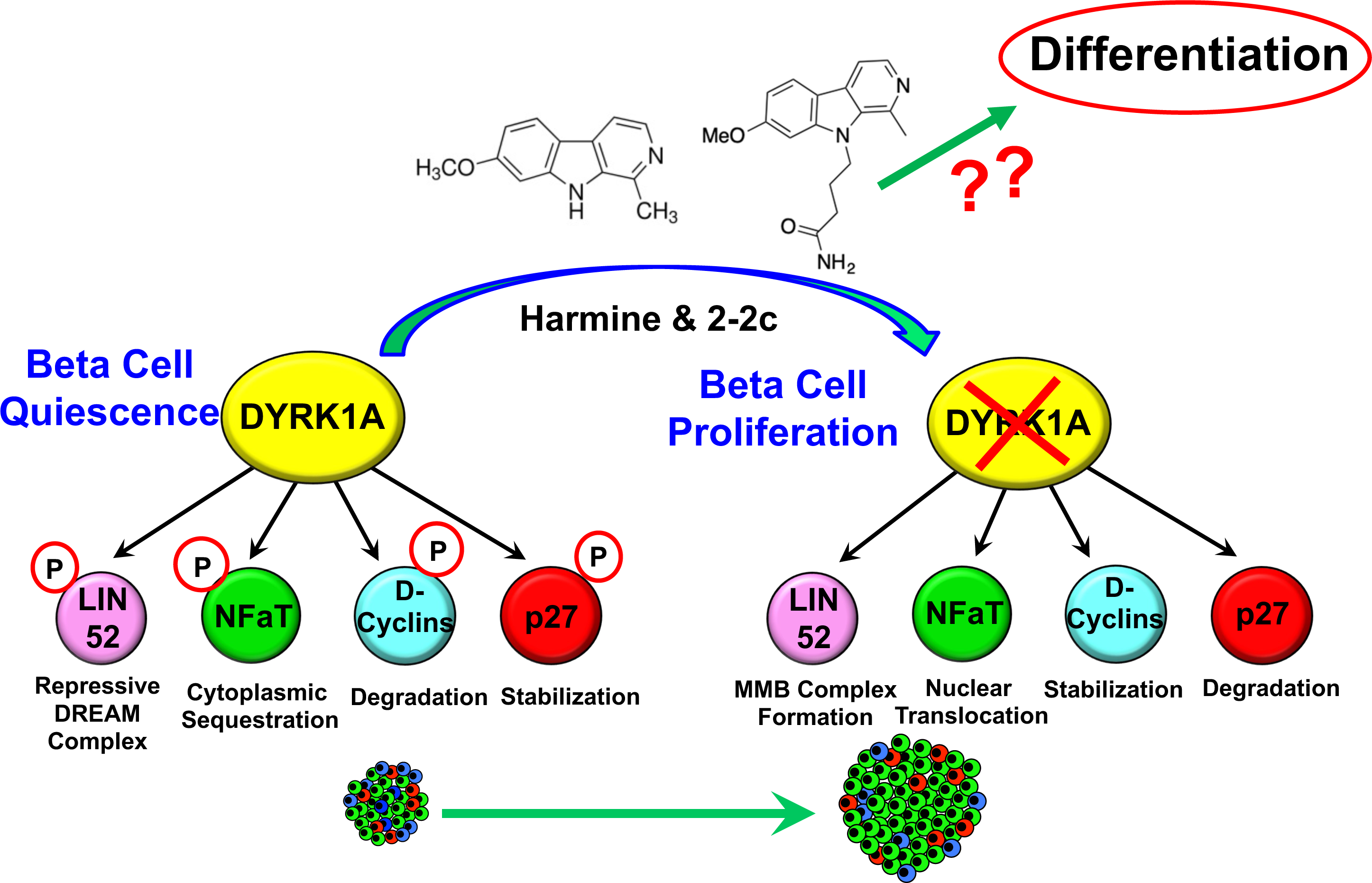
Pictorial Illustration Of The Distinct Mitogenic and Differentiation Pathways Regulated By Harmine, 5-IT, and 2-2c. See text for details

We first selected GNF4877 to study in more detail because, although it is an effective DYRK1A inhibitor, it is highly non-selective (15,24). We speculated that this undesirable non-selectivity might inhibit multiple kinases or other beta cell factors, thereby interfering with its ability to promote differentiation. This proved to be accurate, at least in part, since GNF4877 interferes with the ability of harmine to drive differentiation (**Fig. 5**). Notably, 5-IT is also poorly selective (**Suppl. Fig. 1**)(15,21), yet also effective at enhancing differentiation. This may suggest that although it is promiscuous, it may not interfere with specific kinases or other targets required for differentiation.

As in prior studies (12), while the effects on some of the targets studied here are quantitatively large, e.g., PDX1, NKX6.1, MAFA, MAFB, SLC2A2, PCSK1, and ENTPD3, the effects on INS expression are highly variable and at times small, as illustrated in **Figs. 2**,**5**,**7**,**8**. Nevertheless, in all studies, harmine activates GSIS *in vitro* and *in vivo* and reverses diabetes *in vivo* (10,12,16,17). We interpret the *in vitro* results to reflect the well documented variability among human islet organ donors (34), and the likelihood that while large, rapid changes in transcription factor expression may be required to drive gene expression, small increments in insulin secretion may be sufficient to reverse diabetes.

These observations prompted a candidate-based search for signaling pathways that support human beta cell differentiation and function that might be enhanced by harmine. Since harmine interacts directly with DYRK1A, the obvious mechanistic mechanism for enhanced beta cell differentiation might be loss of DYRK1A function. We and others have found that silencing DYRK1A in human islets drives beta cell proliferation (10,12,13,21), yet we found here that silencing DYRK1A had no effect on beta cell differentiation markers, thus eliminating DYRK1A itself as a pro-differentiation target. We next explored a series of additional candidates implicated in beta cell proliferation, including the CLK family, other DYRKs1B,2,3,4, the DREAM complex, and the NFAT family, all of which are well documented targets of harmine, 2-2c and 5-IT. While silencing DYR1A induces human beta cell proliferation, silencing DYRK1A and each of these other candidate targets had no effect on differentiation marker or transcription factor expression (**Figs. 6,7**). These results leave unanswered the question, “what is the ‘pro-differentiation target’ of harmine, 2-2c, 5-IT (**Fig. 10**)?”. Since a candidate approach was not fruitful, identification of the ultimate target will require an unbiased approach, for example, immunoprecipitation followed by proteome-wide interrogation.

We have shown previously that the proliferative and pro-differentiation effects of harmine extend to islets from human donors with T2D (12). Here, we report that this beneficial effect in T2D islets also applies to 5-IT and 2-2c as well (**Fig. 8**). We also observed that harmine and all of the other small molecule DYRK1A inhibitors reduce expression of ALDH1A3 and TIXNIP, both functional markers and mediators of de-differentiated islets in people with T2D. This supports the concept that DYRK1A inhibitors not only drive proliferation, but may have the ability to reverse established beta cell failure in people with T2D. In contrast to results with the other differentiation markers studied where responses were observed in response exclusively to harmine, 2-2c and 5-IT, the ability to attenuate ALDH31A and TXNIP expression appears to be a common feature shared by all DYRK1A inhibitors. Finally, we observed that harmine limits or reverses the de-differentiating effects of standard glucolipotoxicity and cytokine-induced de-differentiation paradigms.

Overall, these studies further refine the therapeutic utility of the small molecule DYRK1A inhibitor class for beta cell mass expansion and function for T1D and T2D. Notably, these unexpected findings provide an important additional benchmark for assessing and defining the efficacy of human beta cell regenerative drugs: not only must they expand beta cell numbers, the new beta cells must also function at a high level. Importantly, although the mechanisms underlying human beta cell proliferative responses to harmine and other small molecule DYRK1A inhibitor molecules are reasonably clear, despite extensive efforts to eliminate candidate mediators as described herein, the mechanisms underlying the pro-differentiation effects of harmine, 2-2c and 5-IT remain entirely unknown, and require additional study.

## Materials and Methods

### Human Islets

Isolated de-identified human pancreatic islets from otherwise normal organ donors were provided by the NIH Integrated Islet Distribution Program (IIDP, https://iidp.coh.org), Prodo Laboratories, The Alberta Diabetes Institute, and the Transplant Surgery Department, University of Chicago. Details and demographics of the donors and islet preparations are provided in **Suppl. Table 1**. Donor ages ranged from 20 to 68 years old. The mean age (±SEM) was 45.9 ± 13.6 years; mean BMI (±SEM) was 28.8 ± 5.5 kg/m^2^ (range 10-47.6); 35 of 46 were male; 23 were White, 8 Hispanic, 5 Asian, 8 Black, and 2 were not identified; mean cold ischemia time was 469.78 ± 182.23 minutes (range 76–796 minutes); and islet purity ranged from 75% to 99% (mean ± SEM 86.04% ± 5.68%).

### Chemicals

Accutase (Mediatech: 25-058-CL), harmine (286044, Sigma), leucettine-41 (MR-C0023, Adipogen), INDY (4997, Tocris Biosciences), 5-Iodotubercidin (5-IT) (1745, Tocris), Recombinant human TNF-α (210-TA-010, R&D Systems), GNF4877 (HY-129492, MCE), CC-401 (HY-13022A, MCE), 2-2c (synthesized in The Drug Discovery Institute and The Department of Pharmacological Sciences, The Icahn School of Medicine at Mount Sinai).

### Adenoviruses and Transduction

For adenoviral silencing, four target shRNA sequences for each target gene were designed using the Thermo Fisher RNAidesigner online tool. Target sequences were cloned into block-iT U6 RNAi entry vector (K494500, Invitrogen), and silencing efficiency was evaluated in HEK293 cells. The most effective target sequences were used to generate Ad-shRNAs in the Block-iT adenoviral RNAi vector (K494100, Invitrogen). The virus that silences both DYRK1A and DYRK1B simultaneously has been reported previously (13). Adenoviruses for silencing RBL2/p130/RBL1/P107/Rb1, E2F4/ E2F5, and CLK1/CLK2/CLK4 have been reported previously (13). Ad.NFATC1 has been previously described (10). Mouse Ad.ca-nfatc1 and ca-nfatc2 were gifts from Drs. Alan Attie and Mark Keller, Department of Biochemistry, University of Wisconsin. NFAT4 (NFATC3), CA-NFAT2 (CA-NFATC1), CA-NFAT3 (CA-NFATC4) are described in detail in reference (14).

### Human Islet Dispersion and Virus Infection

Islets were centrifuged at 1500 rpm for 5 min, washed twice in phosphate-buffered saline (PBS), re-suspended in 1 ml of Accutase and incubated for 10 min at 37°C. During digestion, the islets were dispersed by gentle pipetting up and down every 5 min for 10 sec. Complete RPMI medium containing 11 mmol/L glucose, 1% penicillin/streptomycin with 10% fetal bovine serum (FBS) was then added to stop the digestion. Dispersed cells were then centrifuged for 5 min at 1500rpm, the supernatants removed, the pellets re-suspended in complete medium, and the cells then plated on coverslips with 30 µl of cell suspension per coverslip. Poly-D-Lysine/Laminin-treated cover slides or chamber slides were used. Cells were allowed to attach for 2 hr at 37°C or were transduced with adenovirus for 2 hr. After 2 hr, 500 µl complete RPMI medium was added in each well to terminate adenoviral transduction. Cells were cultured for 24-96 hr as described in the Figure Legends. Detailed protocols are provided in reference (35).

### Compound Treatments

For compound treatments, dispersed islet cells were allowed to recover on coverslips for 24 hr. RPMI complete medium was then replaced with fresh medium containing harmine or vehicle (0.1% DMSO) for 72-96 hr. Doses used were derived from publications that demonstrated proliferation in human beta cells (10,12,22,23), and were harmine (10 μM), 5-IT (1 μM), GNF4877 (2 μM), INDY (15 μM), Leucettine-41 (20 μM), and CC-401 (10 μM). For the combination treatment with adenovirus and compounds, islet cells were transduced first and then 24 hours later the cells were treated with compound. For the glucolipotoxicity and cytokine studies, islets were exposed to 20 mM glucose and 200 µM palmitate or to 1000 IU/ml TNFα for 72 hours.

### Immunocytochemistry and Antisera

Immunocytochemistry was performed on coverslips fixed in fresh 4% paraformaldehyde. Accutase-dispersed human islet cells were plated on coverslips as previously described (35). Primary antisera were: C-peptide (DSHB GN-ID4); PDX1 (Ab47308, Abcam), NKX6.1 (DSHB, F55A10-c), ENTPD3 (provided by Dr. Alvin Powers, Vanderbilt University).

### qPCR Methods

RNA was isolated and quantitative RT-PCR was performed as described previously (3). Gene expression in dispersed islets was analyzed by real-time PCR performed on a QuantStudio System. Primers are listed in **Suppl. Table 2**.

### Statistics

Statistical analyses were performed using 2-tailed Student’s paired *t-*test as described in the Figure Legends. p values less than 0.05 were considered to be significant.

## Acknowledgments

The authors wish to thank Bonnie and Joel Bergstein, Lonnie and Thomas Schwartz, and Martha and Fred Farkouh families for their constant support of this research. We also thank the NIDDK-supported Human Islet and Adenovirus Core (HIAC) of the Einstein-Sinai Diabetes Research Center (ES-DRC), and the NIDDK Integrated Islet Distribution Program (IIDP), Prodo Laboratories, Patrick MacDonald at the Alberta Diabetes Institute Islet Core at the University of Alberta in Edmonton (www.bcell.org/adi-isletcore) with the assistance of the Human Organ Procurement and Exchange (HOPE) program, Trillium Gift of Life Network (TGLN), and other Canadian organ procurement organizations. We also thank Dr. Tatsuya Kin at the University of Alberta, Edmonton, Alberta, Canada, Dr. Piotr Witkowski at the University of Chicago, Chicago, Illinois, and Dr. Fouad Kandeel at the City of Hope Medical Center, Duarte, California, for providing human organ donor islets. We thank the Icahn School of Medicine Tissue Biorepository for providing deidentified samples of normal human pancreas. We also thank Mark Keller and Alan Attie at the University of Wisconsin for sharing adenoviruses expressing constitutively active mNFAT1 and mNFAT2. This work was supported by NIH grants P30 DK020541, R01 DK116873, R01 DK116904, R01 DK125285, R01 DK105015, R01 DK129196, DK, R01 GM132129, R35 CA232128, and P01 CA203655.

## Author Contributions

P.W., A.F.S. conceived of the studies. P.W., O.W., L.C., H.L., V.W., A.P. performed experiments.

P.W., L.L., A.F.S. analyzed data. P.W., L.C., and A.F.S. wrote the manuscript. K.K., R.J.D. provided reagents and advice. A.G.O., E.K., D.K.S., reviewed data and provided advice.

## Guarantors

P. W. and A.F.S. guarantee the data in this manuscript

## Supplementary Figures and Tables

### Supplementary Figure

**Supplementary Figure 1.**
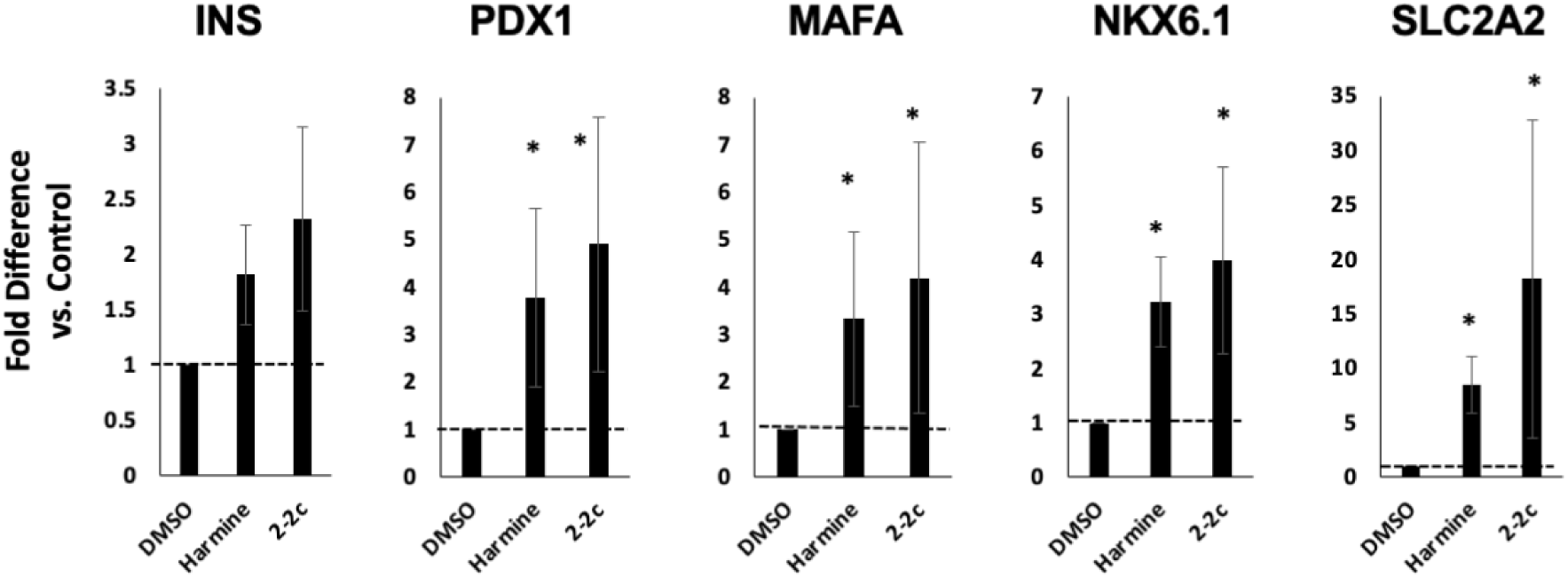
2-2c, Most Potent, Selective and CNS-Avoidant DYRK1A Inhibitor, Mimics Pro-Differentiation Effects of Harmine. Dispersed human islets were treated with DMSO, 2-2c (3µM) or harmine (10µM) for 96 hours. The expression of insulin and other major beta cell transcriptional factors and differentiation markers was assessed by qPCR. *P < 0.05 and **P<0.01 versus control by paired two-tailed t-test. n=5 human islet donors.

### Supplementary Tables

**Supplementary Table 1.** Human Islet Donor Demographics and Characteristics. See .xls File Attached.

| Suppl. Table 1. Islet Donor Demographics |  |  |  |  |  |  |  |  |  |  |
| --- | --- | --- | --- | --- | --- | --- | --- | --- | --- | --- |
| RRID: | Source ID* | Source | Age | M/F | BMI | Ischemia Time (Minutes) | Purity | HbA1c | Race | Cause of Death |
| **Normal Donors** |  |  |  |  |  |  |  |  |  |  |
|  | Hu1197 | ornia Islet Cell Res | 44 | M | 28.84 | 433 | 80 | 5.5 | Hispanic | Stroke |
|  | HP20220 | PRODO | 60 | M | 29.3 | np | 85 | 5.6 | Asian | Stroke |
|  | H268 | U CHICAGO | 24 | F | 39.38 | 697 | np | 5.7 | np | np |
| Samn13254972 |  | IIDP | 60 | F | 44.4 | 374 | 99 | 5.6 | White | Head trauma |
| Samn13149867 |  | IIDP | 28 | M | 30.8 | 503 | 80 | 5.2 | White | Head trauma |
| Samn37737368 |  | IIDP | 50 | M | 38.6 | 362 | 80 | 5.8 | White | Head traum |
|  | HP23270 | PRODO | 34 | M | 27 | np | 85 | 5.5 | Asian | Stroke |
|  | Hp23262 | PRODO | 64 | M | 23.2 | np | 90 | 4.8 | White | Stroke |
|  | HP23258 | PRODO | 39 | M | 26.9 | np | 90 | 5.2 | White | Stroke |
| Samn37350251 |  | IIDP | 43 | F | 29.89 | 300 | 88 | 5.2 | White | Head trauma |
|  | HP23254 | PRODO | 42 | M | 26.5 | np | 85 | 4.8 | Hispanic | Stroke |
| Samn37068473 |  | IIDP | 51 | M | 30.4 | 717 | 80 | 6.3 | White | Anoxia |
|  | Hp23200 | PRODO | 57 | M | 30.8 | np | 95 | 5.2 | Black | Anoxia |
| Samn36471214 |  | IIDP | 40 | M | 24.2 | 300 | 73 | 5.2 | White | Stroke |
|  | HP23189 | PRODO | 33 | F | 28.8 | np | 85 | 5.5 | Asian | Anoxia |
| Samn35848421 |  | IIDP | 22 | M | 31.79 | 540 | 90 | 5.1 | Black | Anoxia |
| Samn35301006 |  | IIDP | 52 | M | 31.6 | 681 | 85 | 4.4 | White | Stroke |
|  | HP23123 | PRODO | 47 | M | 28.3 | np | 90 | 5.7 | Black | Anoxia |
| Samn34411471 |  | IIDP | 45 | M | 33.1 | 360 | 95 | 5.2 | White | Anoxia |
|  | HP23104 | PRODO | 64 | F | 21.9 | np | 95 | 5.8 | White | Stroke |
| Samn34033792 |  | IIDP | 39 | M | 33.4 | 76 | 80 | 5 | White | Head trauma |
|  | HP23019 | PRODO | 55 | M | 30.4 | np | 90 | 5.6 | Hispanic | Anoxia |
| Samn32875450 |  | IIDP | 60 | F | 25.2 | 310 | 75 | 5.7 | White | Stroke |
| Samn32641506 |  | IIDP | 16 | M | 29.5 | 372 | 80 | 5.4 | Hispanic | Head trauma |
|  | HP22362 | PRODO | 51 | M | 24.9 | np | 85 | 5.3 | White | Stroke |
|  | HP22310 | PRODO | 68 | M | 30.7 | np | 90 | 5.4 | White | Anoxia |
| Samn29657580 |  | IIDP | 40 | M | 23 | 692 | 90 | 5.2 | White | Head trauma |
|  | HP22169 | PRODO | 46 | M | 30 | np | 85 | 5.6 | Black | Head trauma |
| Samn29494177 |  | IIDP | 18 | M | 25.3 | 481 | 80 | 5 | Black | Head trauma |
|  | HP18330 | PRODO | 53 | F | 26.3 | np | 90 | 5.4 | White | Anoxia |
|  | HP22105 | PRODO | 58 | F | 28 | np | 90 | 5.1 | White | Head trauma |
| Samn27619977 |  | IIDP | 37 | M | 31.3 | 383 | 90 | np | Hispanic | Head trauma |
|  | HP22095 | PRODO | 33 | M | 25.9 | np | 90 | 5.4 | White | Head trauma |
|  | HP22089 | PRODO | 48 | M | 19 | np | 87 | 5.3 | Hispanic | Stroke |
|  | HP22076 | PRODO | 36 | M | 23.2 | np | 90 | 5.3 | White | Anoxia |
|  | HP22065 | PRODO | 61 | M | 22.4 | np | 90 | 5.2 | Black | Anoxia |
|  | HP22068 | PRODO | 20 | M | 20.8 | np | 90 | 4.6 | Black | Head trauma |
|  | HP22052 | PRODO | 52 | M | 25.4 | np | 90 | 4.2 | White | CVA |
|  | H253 | U CHICAGO | 31 | M | 33 | np | 85 | 5.7 | Hispanic | Anoxia |
| **T2D Donors** |  |  |  |  |  |  |  |  |  |  |
|  | HP21129 | PRODO | 59 | M | 30.4 | np | 80 | 6.7 | Hispanic | Stroke |
|  | HP22044 | PRODO | 55 | M | 30.7 | np | 85 | 6.5 | Asian | Stroke |
|  | HP21182 | PRODO | 62 | F | 20.6 | np | 90 | 5.8 | Asian | Stroke |
|  | HP21263 | PRODO | 62 | M | 34.6 | np | 85 | 6.8 | White | Stroke |
|  | R432 | Edmonton | 66 | F | 36.5 | np | 75 | 8.6 | np | np |
| Samn31065195 |  | IIDP | 48 | M | 23.29 | 796 | 80 | 9.4 | White | Stroke |
| Samn17209843 |  | IIDP | 38 | F | 43.9 | 439 | 80 | 7 | Black | Anoxia |
| np = Not Provided |  |  |  |  |  |  |  |  |  |  |

**Suppl. Table 2.**
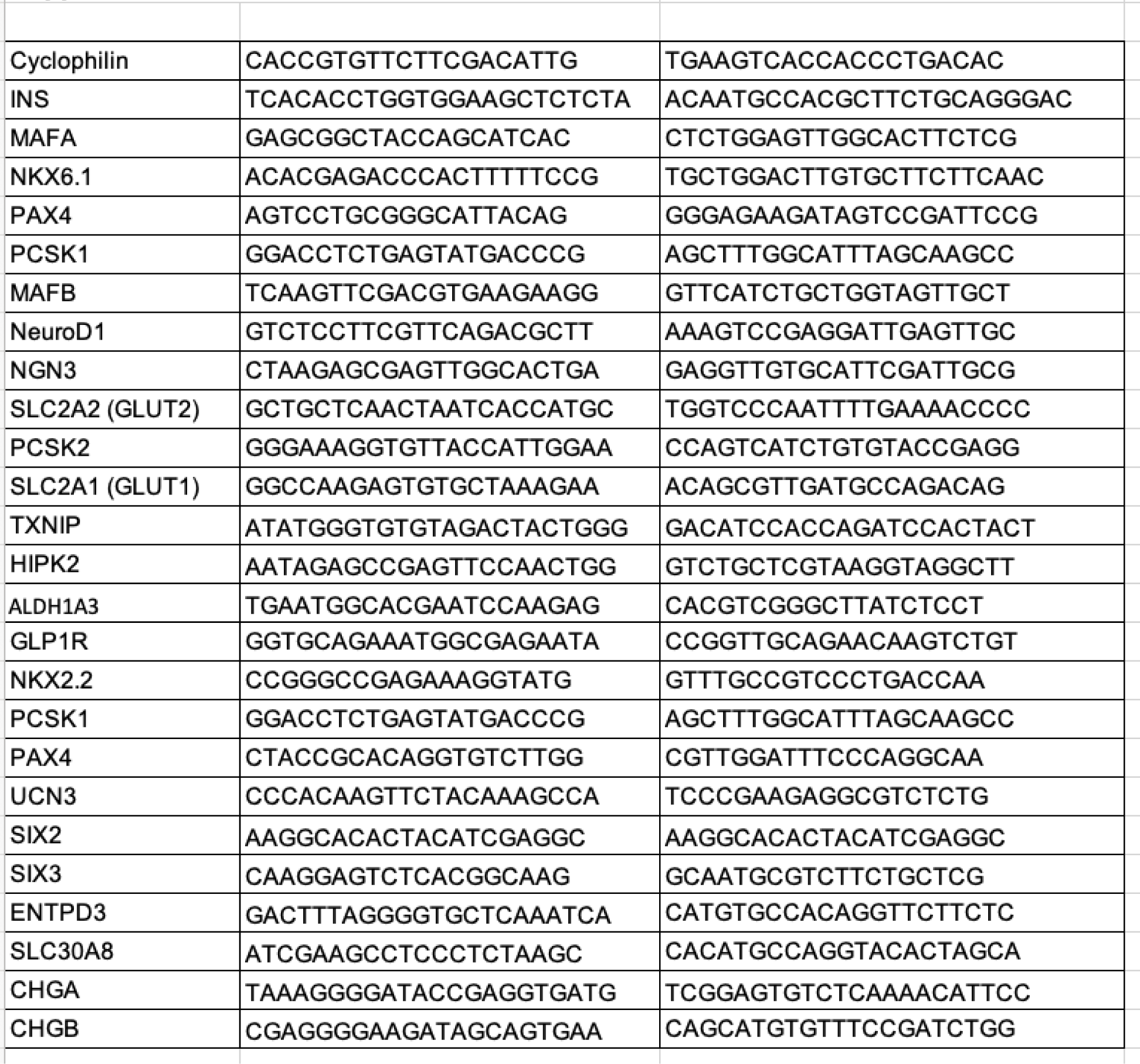
PCR Primers.

